# Illumination mediates a switch in both active sensing and control in weakly electric fish

**DOI:** 10.1101/2024.10.15.618597

**Authors:** Huanying Yeh, Yu Yang, Debojyoti Biswas, Noah J. Cowan

## Abstract

To execute sensory-guided behavior, the nervous system must manage uncertainty within multiple streams of information. There are two highly nonlinear mechanisms for achieving this: 1) sensory reweighting, an *internal neural computation* which places more emphasis on sensory information that exhibits the least uncertainty, e.g. in a Bayesian framework, and 2) active sensing, an *overt behavior* that seeks to improve the quality of sensory information before it enters the nervous system. Here we show that animals solve both of these nonlinear problems concurrently. We studied how the weakly electric glass knifefish *Eigenmannia virescens* alters its movement dynamics under parametric manipulations of illumination. We hypothesized a concomitant switch in both overt active sensing and internal multisensory reweighting. To test this, we varied illumination levels from 0.1 to 210 lx as fish tracked a moving refuge. We discovered that in a neighborhood of a critical threshold (on the order of 1 to 10 lx), small increases in illumination led to dramatic changes in both active sensing and multisensory control, specifically in 1) steep reductions in fish head and tail movements and 2) decreased refuge tracking phase lag. Outside of this threshold, large changes in illumination only caused small changes in active sensing and control. A control-theoretic model that dynamically modulates the weights of vision and electrosense due to illumination changes corroborates our findings. These findings underscore the complex, multipartite, nonlinear nature of locomotor control and the remarkable ability of the nervous system to execute multiple parallel strategies for managing sensory uncertainty.

## Introduction

When navigating the natural environment, animals gather information from various senses, such as visual, tactile, and auditory cues [1–8]. During such navigational behaviors, animals also *internally reweight* streams of sensory information so that larger sensorimotor gains are applied to the sensory streams with the least uncertainties [9–12]. For instance, humans adjust the relative weights of vision, proprioception, and vestibular sensory information to maintain balance under different scenarios, such as when their eyes are closed or when the proprioception of their legs are perturbed [13, 14]. Likewise, electric fish can reweight streams of visual and electrosensory information [15].

Critically, uncertainties in sensing do not only lead to a passive, internal reweighting of sensory information streams; they can also lead to dramatic changes in *overt active sensing behavior*, where animals induce ancillary movements to improve sensory perception in one or more modalities [2,4,16–19]. For example, when crepuscular hawkmoths search for nectar under dim light, they use their proboscis to actively probe a flower’s surface to sense its shape and texture. Here, the temporal dynamics of the active sensing behavior depends on the salience of mechanical features of the flower [20]. While station-keeping inside a refuge, electric fish actively swim back and forth to enforce non-zero relative movement between their receptors and the refuge [21]; these active sensing movements maintain the variance of the sensory error signal in a manner that resembles stochastic resonance [22] but at the behavioral level [23]. Rapid active movements are thought to enhance sensing [24, 25] and feedback control [21,26]. In addition to fore-aft oscillations, these fish also actively bend their tails in response to changes in sensory conditions [17], likely another form of active sensation. Computational models based on information theory [19, 27], as well as control theory and state estimation [28–32], has emerged recently that may help explain the complex, nonlinear dynamics of these active movements.

Active sensing (an overt behavior) and multisensory reweighting (an internal computation) are not mutually exclusive. Thus, we take the next step of investigating how these processes are modulated simultaneously when the available sensory information is perturbed, using the weakly electric glass knifefish *Eigenmannia virescens. E. virescens* relies heavily on its two exteroceptive senses, vision and electrosense [15, 33, 34], to navigate its environment. These fish also excel at a locomotor tracking behavior: by actuating the anal ribbon fin [35,36], they swim forward and backward nearly equally well. Without prior training, these fish readily hover inside a moving refuge with precise positional and temporal coordination (Fig. 1A) [37]. The robust tracking performance, as well as illumination-dependent movement strategies [17, 21], has made *E. virescens* an ideally suited model organism for sensorimotor control studies [38, 39].

**Fig. 1.**
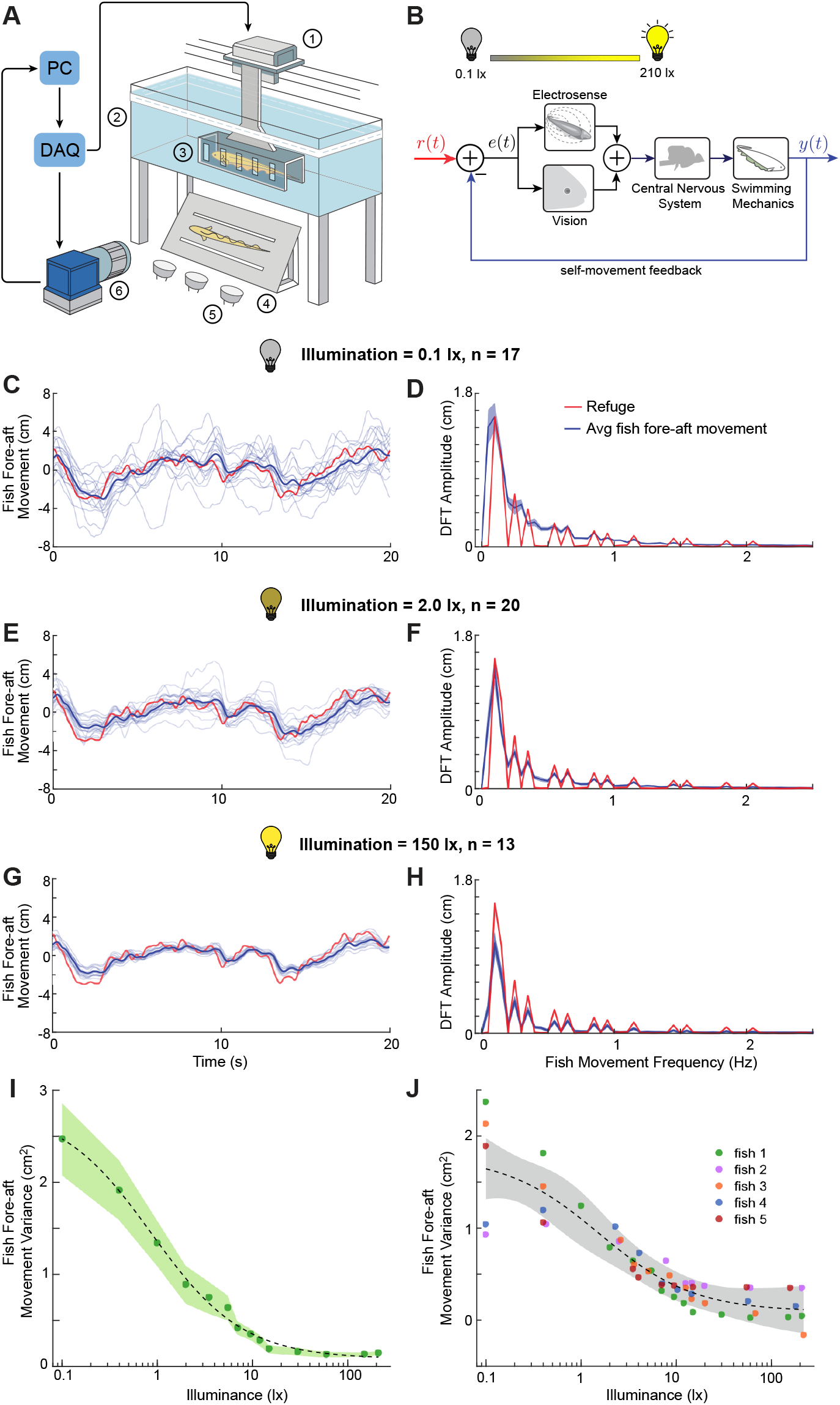
Illumination-dependent behavioral transitions in fish fore-aft movements. **(A)** Schematic of the experimental setup with the following components: 1) linear actuator, 2) LED strip, 3) refuge, 4) first surface mirror, refuge movement *r*(*t*) and the fish’s own movement *y*(*t*). The central nervous system processes the sensory slip and fish. The refuge is actuated with a pre-designed sum-of-sines reference stimulus, *r*(*t*), along the rostrocaudal axis. To 5) infrared (IR) lights, and 6) high-speed camera. **(B)** Block diagram of the closed-loop refuge-tracking system in track the refuge, fish use vision and electrosense to integrate the “sensory slip” *e(t)*, which is the difference between the fore-aft movement (head movement along the rostrocaudal axis) traces (C,E,G) under low (0.1 lx, *n* = 17), medium (2.0 generates the necessary actuation to control swimming mechanics. Experimental manipulation of lighting conditions affects the sensory processing of the sensory slip, which in turn influences the refuge-tracking behavior. **(C-H)** Fish lx, *n =*20) and high (150 lx, *n=* 13) illumination, and the corresponding discrete Fourier Transform (DFT) amplitudes (D,F,H) from a single representative fish (blue) tracking the refuge movement (red). Lighter shades in (C,E,G): individual position traces. Darker shades in (C,E,G): average fish fore-aft movements. The solid lines in (D,F,H) denote the mean values of all trials’ DFT amplitudes. The shaded regions in (D,F,H) show the standard error of mean (s.e.m). **(I)** Fish fore-aft movement variance of the same fish vs. illumination. Each marker represents the mean across trials, while the shaded region denotes the s.e.m. The dashed black line denotes the sigmoidal fit to the data. **(J)** Combined fish fore-aft movement variance for all individuals (*N =*5) vs. illumination. Different colors represent different fish; each marker is the mean of 8–20 trials. The dashed black line and the gray-shaded region denote the sigmoidal fit to the combined data and 95% prediction interval, respectively. See Table S2 and Fig. S1B,C for fitting details.

In this study, we parametrically examined the effect of illumination on both active sensing and multisensory control as *E. virescens* tracked a pseudo-randomly moving refuge under a broad range of lighting conditions (0.1 to 210 lx). We discovered that illumination mediated a switch at a threshold (on the order of 1 to 10 lx), in which slight increases or decreases in illumination produced large decreases or increases, respectively, in active sensing movements. We also discovered a similar threshold in the fish refuge tracking dynamics, especially the phase lags which rose sharply at frequencies above 1 Hz when illumination was decreased near the threshold. We formulated a control-theoretic multisensory reweighting model that recapitulates our findings, indicating that *E. virescens* dynamically changed the relative sensory weights between electrosense and vision. This mechanism appears similar to how hawkmoths dynamically reweight visual and mechanosensory information when feeding from a moving flower [40].

The illumination threshold we discovered, which simultaneously and abruptly alters both internal sensory reweighting and overt active sensing behavior, could foster further investigation on how animals respond to sensory conflicts [15] under different sensory weights, and on how animals solve the explore-vs.-exploit problem [28] under a range of sensory conditions.

## Results

We recorded tracking responses of untethered fish inside a moving refuge using a camera that imaged the fish from below (Fig. 1A). Then, we digitized the horizontal position and 2D body shape of five individual fish in 20 s trial duration for 8–20 trials per condition. Experiments were performed for illumination levels ranging from 0.1 to 210 lx on a logarithmic scale (Table S1). This illumination range, from a dark to a well-lit aquarium, enabled us to test the parametric effects of illumination on active sensing and locomotor control during refuge tracking (Fig. 1B). Lighting levels were randomized across trials. The refuge movement followed a pseudo-random sum-of-sines stimulus comprising 12 frequency components (see Material and Methods) [41, 42].

### Fish exhibit an illumination-dependent switch in active sensing

Across illumination levels, the dominant frequency peaks in fish fore-aft movement (*head* movement along the *rostrocaudal* axis) aligned with those of the refuge (Fig. 1D,F,H). The averaged movement traces of fish and refuge were highly correlated (Fig. S1A). Together, this implies that fish consistently tracked the refuge across all tested illumination levels.

We observed an increase in the variability in fish fore-aft movements under decreasing illumination (see progression in Fig. 1C,E,G). The fore-aft movements were composed of the refuge-tracking movements (sharp peaks in the frequency response at the 12 refuge stimulus frequencies) as well as ancillary active sensing movement which can be observed most clearly as highly variable movement in the time domain (Fig. 1C,E,G).

The fore-aft movement variance as a function of illumination from one representative fish (Fig. 1I) and that from the population (Fig. 1J), indicated an increase in fore-aft active sensing movements as a function of decreased sensory salience, as expected from prior literature [17, 21, 28]. Building on prior work, we examined active sensing in relation to parametrically varied illumination levels. Our finding revealed that the variance of fish fore-aft movement changed in a sigmoidal pattern [43–45], suggesting the presence of an illumination-mediated switch. To fully characterize the transition, we fitted a sigmoidal function to the movement variance as a function of the log of the illumination level (see Material and Methods and Table S2). The sigmoidal fit was also better than polynomial fits, as evidenced by both Adjusted *R*^2^ and the Akaike Information Criterion (AIC), shown in Fig. S1B,C. The sigmoidal fit of the combined fish data indicated the presence of an illumination-dependent threshold. At 0.95 lx the fish fore-aft movement variance began to decrease sharply with respect to the illumination, until reaching 5.32 lx (Fig. 1J, Table S2). Outside this illumination range, the rate of change in variance became largely insensitive to illumination changes.

In addition to fore-aft active sensing movements, *E. virescens* also bends its body and tail to enhance active sensing [17]. Here, we analyzed whole-body bending motion as a parametric function of illumination from 0.1 to 210 lx by tracking 12 points along the center line of the fish’s body, from the head tip *b*_0_ to the tail end *b*_11_ (Fig. 2A,B, see Supplemental Methods for spacing methods). We analyzed body bending and tail movements relative to the fish’s body frame by establishing a body frame at the head position *b*_1_, with orientation given by best-fit line through the three points *b*_0_, *b*_1_, and *b*_2_ (see Material and Methods and Fig. S4).

**Fig. 2.**
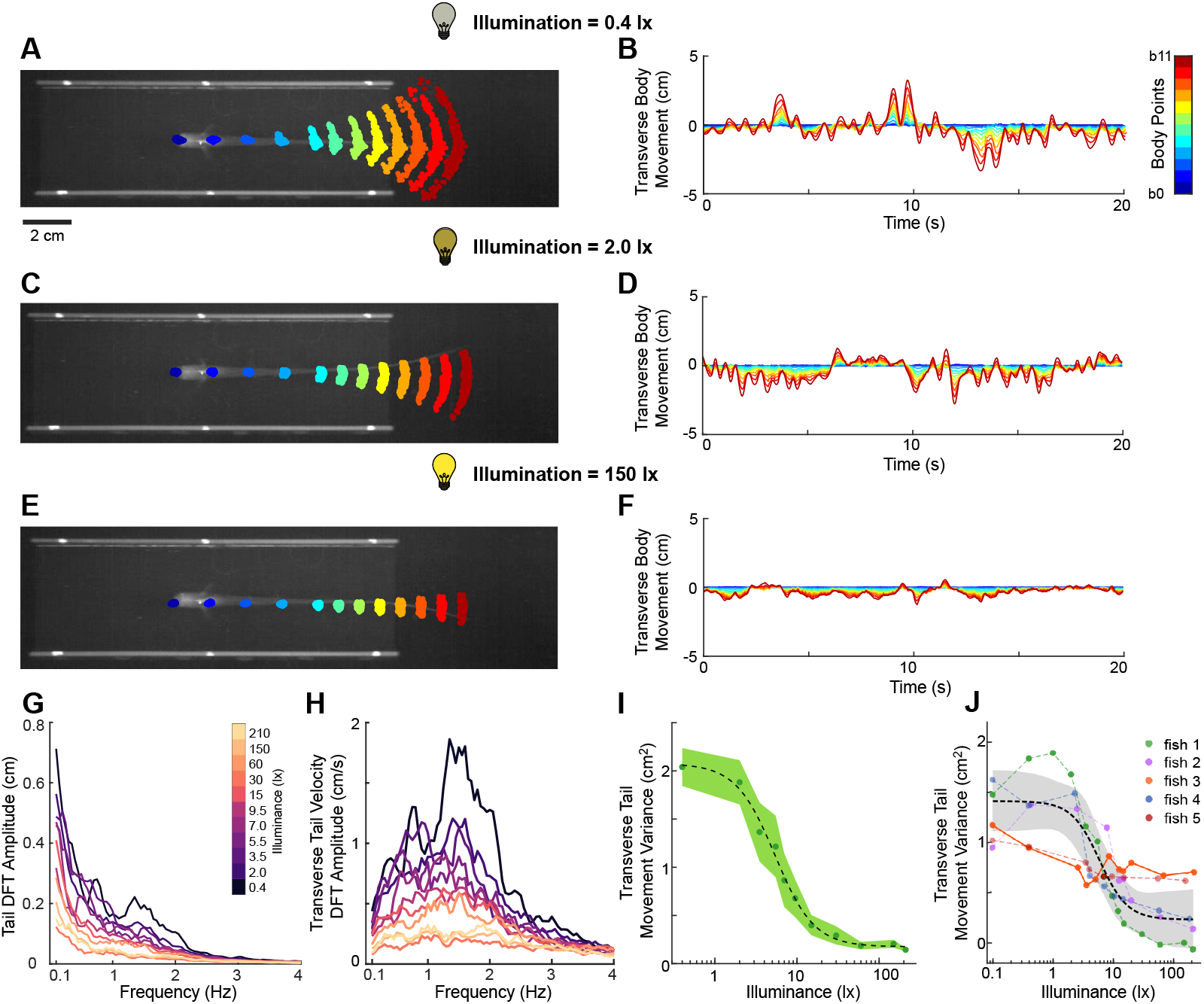
Fish body dynamics changed in response to illumination. **(A-F)** Fish body movements over 20 seconds are depicted by 12 colored markers overlaid on the fish bottom view images (A,C,E). The corresponding temporal traces of the transverse fish body movements in time domain are shown in (B,D,F) under three illuminations (0.4, 2.0 and 150 lx). **(G**,**H)** Average discrete Fourier transform (DFT) amplitude of transverse tail movement (G) and the corresponding velocity (H) across trials from one sampled fish (fish 1). Different colors denote different illumination levels. **(I)** Variance of transverse tail movement across different illumination from one sampled fish (fish 1). The markers and shaded regions denote the mean and the standard error of mean, respectively. The dashed black line denotes the sigmoidal fit to the data. See Table S2 for fitting details. The data corresponding to 0.1 lx, 1 lx and 12 lx are omitted in (G–I) due small sample size (n ≤ 3); see Fig. S1D, Fig. S2 for full version. **(J)** Combined transverse tail movement variance for all to individuals (*N* = 5) under different illumination. Different colors represent different fish; each marker is the mean of 2–17 trials. Except for one of the fish (orange, Adj *R*^2^(sigmoid) = 0.62, Adj *R*^2^(quadratic) = 0.79), all other fish showed a sigmoidal trend (Adj *R*^2^(sigmoid) ≥ 0.9, *N* = 4). The black dashed line and the gray-shaded region denote the sigmoidal fit to the combined data and 95% prediction interval, respectively. See Table S2 for fitting details.

We found that the tail beat amplitude (from transverse movement of *b*_11_) decreased under higher illumination (Fig. 2A–F) as did the corresponding Fourier spectrum amplitude (Fig. 2G, Fig. S2). The Fourier spectrum of velocity revealed a prominent peak between 1.0 and 1.5 Hz that also decreased with increasing illumination (Fig. 2H, Fig. S2). We observed a pronounced illumination-dependence in tail movement, again following a sigmoidal trend (Adj *R*^2^ ≥ 0.9) in four out of five fish (Fig. 2J, trend from a sampled fish shown in Fig. 2I and Fig. S1D). However, one fish’s (orange) decrease in variance was better fit by a quadratic (Adj *R*^2^ = 0.79) than the sigmoidal model (Adj *R*^2^ = 0.62), shown in Fig. 2J. Taken together, these findings demonstrate that fish actively regulate their body movements in response to changing illumination during refuge-tracking tasks in such a way that the intensity of these active sensing movements declines sharply once illumination surpasses a certain threshold.

### Tracking phase lag decreases sharply near the illumination threshold

As fish actively moved for sensing, they simultaneously tracked the trajectories of the refuge, as seen in the time-domain tracking response (Fig. 1C–H) [39]. How does illumination mediate performances during this refuge tracking task? We answered this question by driving the refuge with a sum-of-sines stimulus [41], measuring fish position in response to the refuge motion, and applying system identification approaches to estimate the closed-loop frequency responses of fish refuge tracking and quantify tracking performance over the frequency range of 0.1 to 2.05 Hz [41, 42, 46, 47]. We found that at low frequencies (<0.2 Hz), fish performance approached perfect tracking across illumination conditions, with a gain close to 1 and a phase lag near 0^○^ (Fig. 3A,B). At higher frequencies, the gain and phase “rolled off” to lower values, as expected. Notably, there were no significant differences in gain across illumination levels, but a clear increase in phase (decrease in phase lag) as illumination increased, especially at high frequencies greater than 1.55 Hz. The three highest sum-of-sines frequencies (1.55 Hz, 1.85 Hz, and 2.05 Hz) displayed the most prominent illumination dependence.

**Fig. 3.**
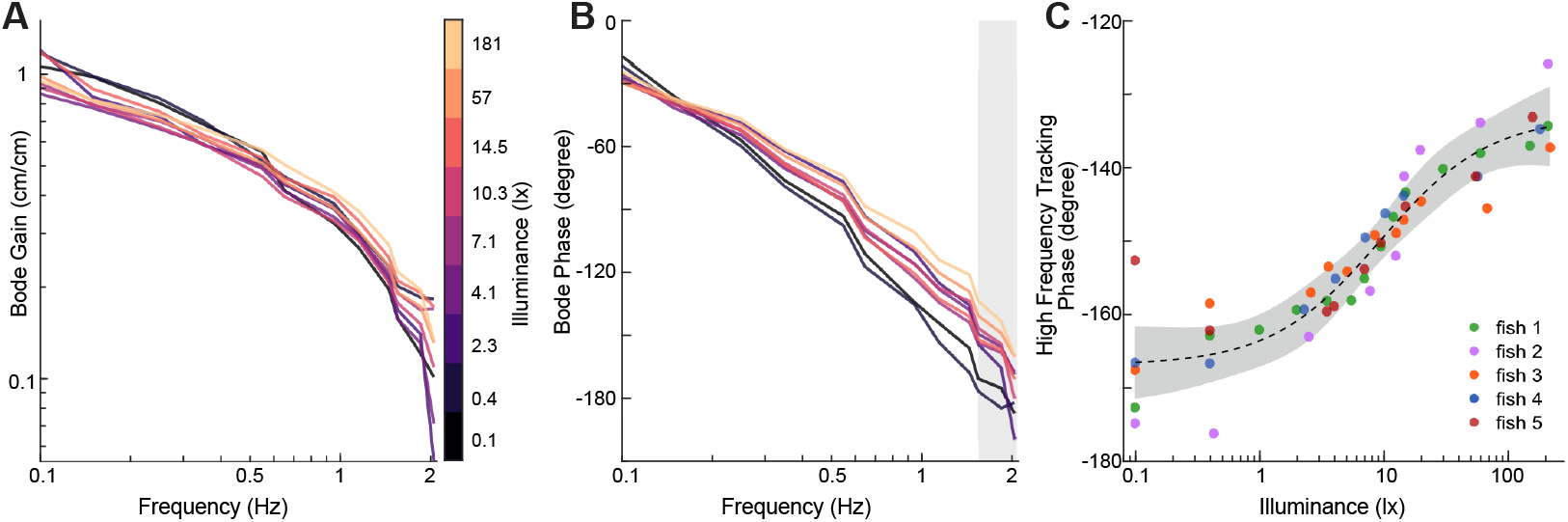
The closed-loop frequency responses of fish tracking. **(A**,**B)** Representative Bode gain (A) and phase (B) plot of fish frequency responses v.s. illumination. Bode gains tightly distribute over the refuge frequency range (0.1 – 2.05 Hz), whereas Bode phase values, initially tightly grouped, become more dispersed at higher frequencies (> 0.5 Hz). Different colors denote the illumination scale. See Fig. S3 for results from all five fish. **(C)** Closed-loop high frequency tracking phase vs. illumination for all individuals (*N* = 5), computed by averaging the phase in the 1.55 – 2.05 Hz frequency range (shaded region in (B)). Each color represents a subject; each marker is the mean of 4 – 17 trials. The dashed black line and the gray-shaded region denote the sigmoidal fit to the combined data and 95% prediction interval, respectively. See Table S2 for details on fitting.

To quantify this trend, we averaged the tracking phase values for these three highest sum-of-sines frequencies and observed a sigmoidal trend in tracking phase (Fig. 3C), which again bears striking similarity to those in active sensing fore-aft (head) movement (Fig. 1I,J) and tail movement (Fig. 2I,J). For all five fish, sigmoidal fits were better than polynomial fits (Table S2). This combined data exhibits a threshold region between 3.56 lx and 18.17 lx. Within this range, phase lag decreased sharply as illumination increased. Outside of this range, the phase lag was generally insensitive to changes in illumination. It is possible that this change in phase lag arises from an illumination dependent delay [48] or perhaps a more complex multisensory reweighting [40], which we examine in the next section.

### Fish dynamically reweight vision and electrosense under changes in illumination

*E. virescens* primarily uses its visual and electrosensory systems to process environmental cues. How does it tune these sensing modalities to bring about the illumination-mediated behavioral transitions? To address this question, we proposed a control-theoretic model involving these two sensory systems. We modeled the dynamics as a closed-loop system (Fig. 1B), including distinct sensory dynamics for vision, *S*_*V*_, and electrosense, *S*_*E*_. In the model, the controller, *C*, represents the sensorimotor transform performed by the nervous system, and the musculoskeletal plant, *P*, represents how the swimming dynamics transforms motor commands into fish movement (Fig. 1B).

Previous work explored the role of illumination on flower-tracking behavior in hawkmoths [48, 49]. In an illumination-dependent control system model, a single temporal processing delay was sufficient to explain the moth flower tracking behavior in different illumination conditions [48,49]. However, we found that simply adding a delay block (Fig. 4A) with the transfer function *e*^−*sτ*(*λ*)^, where *λ* is the illumination level and *τ* (*λ*) is the illumination-dependent time delay, failed to explain the fish behavior (Fig. 4B,C).

**Fig. 4.**
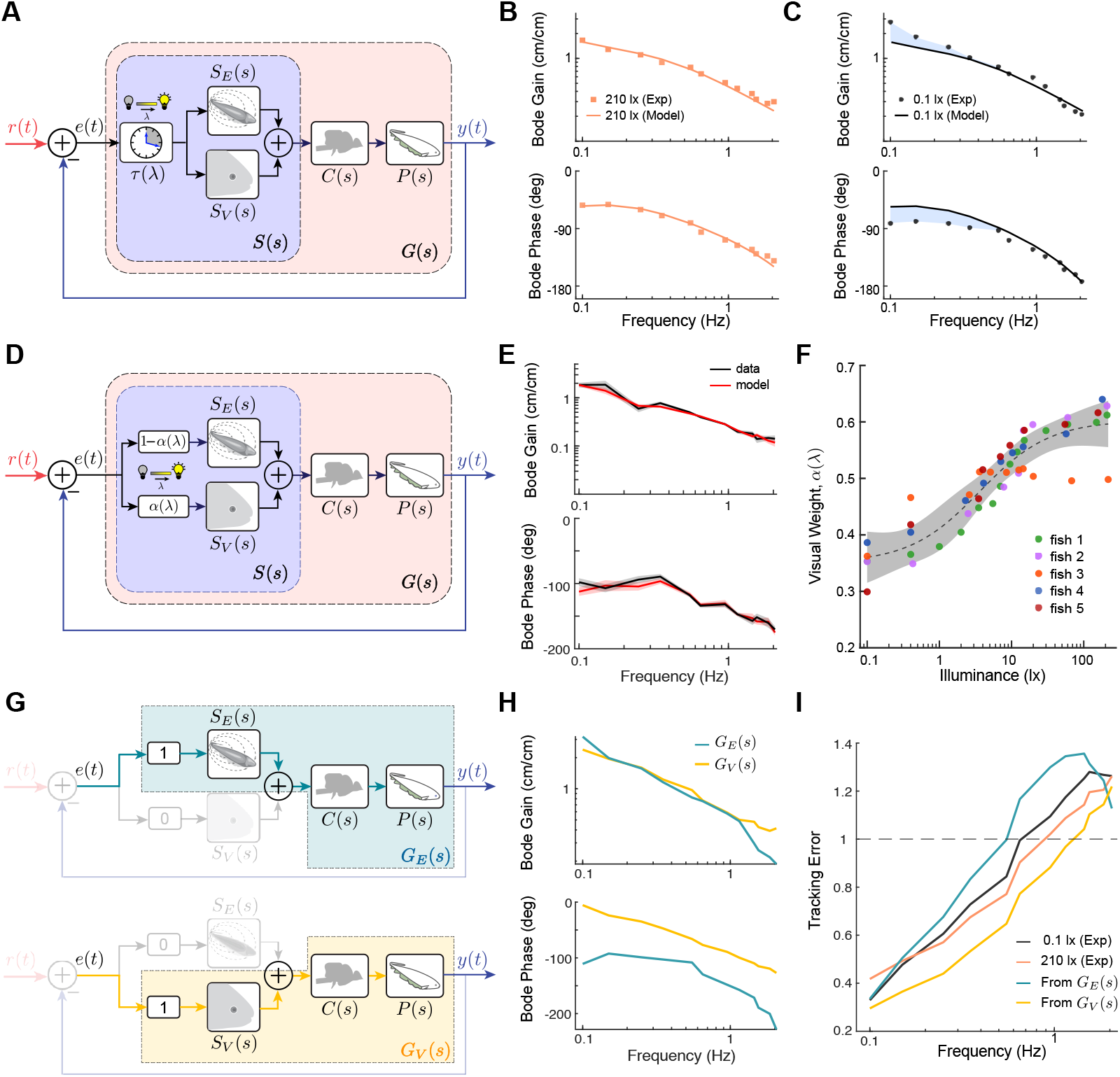
Illumination-dependent sensory reweighting explains the observed behavior in fish. **(A)** A schematic showing the candidate heuristic of illumination-dependent sensory delay. The sensory system is represented by *S* (*s*) (purple dashed box), containing the illumination-dependent sensory delay, *τ* (*λ*), visual system *S*_*V*_ (*s*), and electrosensory system *S*_*E*_(*s*). This heuristic is under the assumption that more sensory delay *τ* is needed as illumination level *λ* decreases. The central nervous system and the swimming dynamics of fish are depicted as the controller, *C*(*s*), and plant, *P* (*s*), respectively. The open-loop sensory system, controller, and plant cascade is depicted as *G* (*s*) (pale pink dashed box). This process is under feedback control. **(B)** Bode plots of open-loop sensory system, controller, and plant cascade *G (s*), generated from the experimental data of a representative fish (fish 1) under high (210 lx, peach squares) illumination condition. The best fit of data in high (210 lx) illumination obtained from the variable delay model is shown in peach colored line. **(C)** Bode plots of *G (s*) generated from the experimental data of the same representative fish from (B) under low (0.1 lx, black circles) illumination condition. Adding a pure delay to the fitting in high (210 lx) illumination in (B) (black line) does not predict data in low (0.1 lx, black circles) illumination condition well, providing the differences in gain and phase at low frequencies below 0.55 Hz (blue shaded region). **(D)** Schematic showing the proposed sensory reweighting model, where “sensory slip” *e (t*) is processed by the sensory system *S (s*) (purple dashed box), containing visual pathway *S*_*V*_ (*s*) and electrosensory pathway *S*_*E*_(*s*) with sensory weights, *α*(*λ*) and 1 − *α*(*λ*), respectively. *α* is a variable depending on illumination level *λ*. **(E)** Bode plot of the averaged *G (s*), from bootstrapped datasets (black) and the corresponding fitted result (red) from the sensory reweighting model for a representative fish (fish 4) at 7.1 lx. The lines and shaded regions denote the corresponding mean and standard deviation of the 100 bootstrapped data and their corresponding fitted results, respectively. **(F)** Sensory weights of the visual pathway, *α*(*λ*) across different illumination levels are presented for all individuals (*N=* 5). Different colors represent different fish. The dashed black line and the gray-shaded region denote the sigmoidal fit to the combined data and 95% prediction interval, respectively. See Table S2 for fitting details. The characteristic “S” shape denotes that above a certain illumination threshold, fish began to rely more on vision, resulting in higher visual weight. **(G)** The hypothetical “electrosense only” pathway (top, deep aquamarine color dashed box) with transfer function *G*_*E*_ (*s*) and “vision only” pathway (bottom, yellow dashed box) with transfer function *G*_*V*_ (*s*). **(H)** Bode gain (top) and phase (bottom) plots for the open-loop sensory system, controller, and plant cascade of a representative fish (fish 1) under the hypothetical “electrosense only” (deep aquamarine) and “vision only” (golden yellow) conditions. In the “vision only” condition, there is a notably lower phase lag and a flatter gain roll-off compared with “electrosense only” condition. **(I)** Frequency vs. frequency domain tracking error plots of the experimental data from a representative fish (fish 1) at 0.1 lx and 210 lx, placed adjacent to the “vision only” and “electrosense only” simulated results from the experimental data for the same fish. The similar trends align with experimental observations, where higher electrosensory weights correspond to higher tracking error, especially at frequencies >0.55 Hz.

The time-domain and frequency-domain analyses presented earlier suggest an illumination-mediated switching behavior in fish. Below a certain illumination value, fish engaged in more ancillary active sensing movements, while above a certain higher illumination value, fish movement variance decreased. At low visual salience conditions, the extraneous movements enhance active sensing with electrosense, whereas in well-lit conditions, such movements attenuated, suggesting a heavier reliance on vision. Based on these observations, we propose an alternative illumination-dependent sensory reweighting model, where the error signal “sensory slip” *e*(*t*) is processed by the visual and electrosensory pathways with illumination-dependent sensory weights, *α*(*λ*) and 1 − *α*(*λ*), respectively, in a manner similar to a model recently proposed for hawkmoths [40]. In this model, we assume that the transfer functions for vision *S*_*V*_ (*s*), electrosense *S*_*E*_ (*s*), the controller *C (s*), and the plant *P (s*) are all illumination-invariant (Fig. 4D). Thus, the impact of illumination is solely captured by the sensory weights, *α (λ*) for vision and 1 − *α (λ*) for electrosense, in which a single illumination-dependent parameter *α* serves to complementarily reweight vision and electrosense.

To test the model, we first computed the open-loop frequency response function (FRF), *G(jω*), from the experimental data. After that, we derived an optimal *G*_*V(*_ *jω*), *G*_*E*_ (*jω*), and a set of *α* for each illumination condition tested which minimized the cumulative mean-squared error (MSE) between model 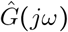 and data *G jω* at frequencies of interests from all illumination levels (see Material and methods and Supplemental Methods for further details). The fitted open-loop FRFs, based on the sensory reweighting model, dramatically improved the fit when compared with the “sensory delay” model (Fig. 4A-C), matching the experimental gain and phase responses (e.g. Fig. 4E). However, the fitting quality declined for low illuminations, due to greater trial-by-trial variance in the data. Interestingly, the computed visual weight *α* presented similar sigmoidal trend across different fish (Fig. 4F) as found for other parameters of active sensing and control.

The proposed sensory reweighting model allows us to compute the open-loop transfer function from error *e*(*t*) to movement *y*(*t*) as follows:

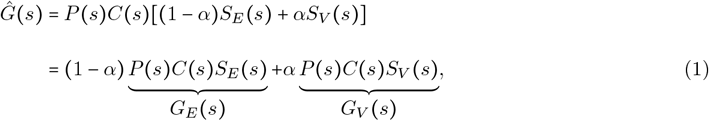

Here, the terms *G*_*E*_ (*s*) and *G*_*V*_ (*s*) can be thought of as the “electrosense only” and “vision only” pathways, respectively (Fig. 4G), as would be found in a hypothetical sensory ablation experiment [50]. We found the illumination-invariant open-loop system transfer function for the “electrosense only” condition *G*_*E*_ (*s*) has greater phase lag and a sharper gain roll-off (Fig. 4H) compared to that of “vision only” *G*_*V*_ (*s*). Moreover, fish exhibited a significantly lower frequency domain tracking error across the frequency range of 0.1 – 2.05 Hz in “vision only” hypothetical condition compared with that in the “electrosense only” hypothetical condition (Fig. 4I). The experimental frequency domain tracking error at the lowest (0.1 lx) and highest (210 lx) illumination conditions were between the two hypothetical extreme conditions (Fig. 4I) as expected. Taken together, these data indicate a complementary illuminance-dependent sensory switch in the weights applied to vision and electrosense.

## Discussion

In this study, we applied a single-parameter change in visual salience to evaluate refuge tracking of weakly electric fish. Under a broad range of illumination levels, we examined the body motion and frequency responses of the fish. At lower illumination levels, fish not only presented higher head-position variance across trials and more vigorous tail-bending movements, but also presented higher phase lag in the high-frequency responses. The former implied more overt active movements for electrosensation, while the latter implied the outcome of an internal reweighting of multisensory control. Notably, we observed a similar illuminance threshold across all metrics, namely trial-by-trial variance in head locations, tail bending, and refuge tracking phase lag. Around this threshold, small changes of illumination level brought about significant, concomitant changes in both nonlinear processes—active sensing and multisensory reweighting. This illumination threshold suggested a switch from an “electrosense dominant” sensing mode to a “vision dominant” mode as the illumination level increased, both in terms of how sensory streams are processed and by how the animal actively gathers information. This research sheds new light on illumination-dependent sensing and control in object tracking tasks, enhancing our understanding of animal active sensing, multi-sensory integration, and these deeply interconnected nonlinear processes that underly locomotor control.

### Active movements enhances sensory processing

We observed that as the lighting intensity decreased, the trial-by-trial variance in fish head movement trajectories and the tail-bending magnitudes both increased. This supports earlier findings that *E. virescens* engages in more vigorous movements to enhance electrosense and compensate for the lack of vision [17,21,51,52] to prey-capture [53–55]. In low light, weakly electric fish usually generate high frequency fore-aft movements [17, 19, 28, 51] and actively bend the trunk and tail [17, 53, 55–57]. Such movements, the hallmarks of active sensing, not only increase the overall sensory volume of the fish [25], but may also serve to “up-modulate” frequency content to match the spatial-temporal properties of neural circuits [17] and prevent perceptual fading [58]. Specifically, tail bending induces changes in the relative geometry between the electrosensory organ and the electroreceptor, which generally causes an amplitude modulation: a stronger electric field on the ipsilateral side of bending while a weaker stimulation on the contralateral side [59, 60]. The principal frequency-domain velocity amplitude of tail bending centered at around 1.0 – 1.5 Hz across illumination levels (Fig. 2H, Fig. S2F–J). Such modulation might be extracted by neurons that are selective for amplitude modulations at the same frequency band in fish mid-brain [61, 62]. Tail bending also affects electrosense by retuning neurophysiological activities; the repeated tail beating could be canceled by descending feedback sent to the Pyramidal cells in the electrosensory lateral line lobe (ELL), a cerebellar-like structure of fish [17, 63–65].

Indeed, whole body fore-aft active movements driven by the ribbon fin [35, 36] and active tail bending [17] likely help fish obtain sensory information. However, at the same time, these motions incur energetic costs [25]. Several theories have been proposed recently for rationalizing these movements in the face of the associated costs [19, 28, 29] and future work could extend these theories to allow for parametric manipulations of salience in one modality (i.e. illumination) to examine which (if any) of the models predict the switches in active sensing and control we observed experimentally.

### Ecological significance of the illumination threshold

The illumination threshold on the order of approximately 1 to 10 lx observed in this study reflects an intricate feature of animal sensorimotor control systems. Sensory movement strategies were rapidly adjusted in response to changes of illumination throughout the threshold, then “saturated” as illumination level surpassed around the order of 10 lx (Fig. 1I,J, Fig. 2I,J). Note that across experimental trials, the illumination conditions were applied in a pseudo-randomized order, so it was unlikely for the fish to adapt to certain patterns of illumination change. Instead, the results suggested that within the neighborhood of illumination threshold, visual receptors of *E. virescens* seem to be more sensitive to illumination change, so that fish can gather adequate quantity of visual information and quickly reweight vision and other senses. The threshold range occurs at relatively low illumination, which could be result from *E. virescens*’ ecological niche; as a nocturnal species living in turbid water and in complex root and littoral habitats in South America, *E. virescens* may have visual receptors more suitable for relatively low light intensities [66, 67].

### Sensory reweighting and integration in sensorimotor control

We proposed a control-theoretic sensory reweighting model for the refuge tracking task of weakly electric fish, which captured responses over a broad range of illumination levels. Under the assumption that fish dynamically integrate visual and electrosensory pathways, our model suggested that as illumination levels decreased, fish tended to adjust the weights of two sensory pathways and put more weight on the electrosensory pathway than the visual one. From Fig. 4H, we see that electrosensory pathway possesses larger phase lag than that in the visual pathway, thus requiring longer integration time than the visual pathway. This was surprising to us and the reasons are not entirely clear. One speculation is that electrosense relies on whole-body active sensing movements, which by their nature, require time to complete. In any case, the increased phase lag in lower illumination appears to be due to the increased weight of electrosense, as well as the intrinsically larger phase lag of the electrosensory pathway.

We applied linear time-invariant (LTI) control theory tools to construct our model, i.e., a linear combination of illumination-independent transfer functions of visual and electrosensory pathways multiplied by their corresponding illumination-dependent weights. It is critical to note, however, that the sensory reweighting process itself is fundamentally nonlinear. Our application of LTI theory inherently assumes that the nonlinear sensory reweighting process was sufficiently *fast* that, during any given trial, the weights for each sensory pathway were time-invariant after a brief reweighting process. This is, in a sense, opposite to the assumption by Yang et al. (2021) [68] and Yang et al. (2024) [42] in which LTI models were used to study learning, wherein the assumption was that the learning rate was sufficiently *slow*. Moreover, the LTI modeling assumes that the highly nonlinear active sensing movements “average out” over multiple trials, revealing dynamics that—on average—appear to be LTI. In essence, LTI modeling has its limitations when applied to nonlinear systems. However, by carefully managing the different timescales of nonlinear processes (such as fast sensory reweighting or slower learning dynamics), we can effectively apply LTI models in a piecewise fashion to help characterize nonlinearities in the controller dynamics.

Sensory reweighting is commonly studied across taxa. For instance, in human balancing control which integrates visual, vestibular, and proprioceptive sensory information, researchers usually apply sensory perturbations in the environments and test the relative contribution that different sensory systems make for completing the task [13, 69, 70]. In cross-modal object recognition of weakly electric fish *Gnathonemus petersii*, fish were trained to use either vision or electrosense to discriminate two objects; they could accomplish the task using only the untrained sense [71]. Results showed that they dynamically reweighted the object-related sensory input based on their reliability to minimize the uncertainty and optimize sensory integration [71]. Researchers also found that rats and humans dynamically adjusted weights assigned to each sensing modality based on its reliability to help decision-making [72]. Theories such as nonlinear adaptive control models [14] and a Bayesian framework [73] have been applied to model the multisensory reweighting.

Researchers also introduce sensory conflicts, i.e., perturbing the system simultaneously with different sensory inputs, to test how different sensing modalities integrate in animal sensorimotor behaviors. Some examples include refuge tracking of weakly electric fish (vision and electrosense) [15], wing steering in fruit flies (vision and haltere mechanosense) [74], and flower tracking of hawkmoths (vision and proboscis mechanosense) [40, 50]. Although neural systems adopt complex encoding/decoding mechanisms possessing nonlinearity [75–78], many studies have found that sensory information integrates approximately linearly under conflicts across different sensorimotor behaviors [15, 40, 50]. Conversely, some examples show that linearity does not hold for sensory integration. For instance, linear superposition of haltere-driven and visually-driven responses of *Drosophila* predicted their wing steering responses but not gaze responses [74]. These studies illustrate that despite the elements in nervous systems are intrinsically nonlinear, at least within a certain range of conditions for behaviors, the combined activity often presents itself linearity [79]. Further investigation at both the behavioral and circuit levels can reveal more insights on how these highly nonlinear processes integrate across various sensing and actuation modalities across animals to achieve robust closed-loop performance.

## Acknowledgments

We thank Eric S. Fortune for helpful discussions and suggestions; we also thank Calvin Yeh for significant contributions to the machine learning-based image processing pipeline. This work was supported by the Office of Naval Research under grant no. N00014-21-1-2431 (N.J.C), the National Science Foundation under grant no. 2011619 (N.J.C), and the Provost’s Undergraduate Research Award (PURA) at Johns Hopkins University (H.Y.).

## Author Contributions

H.Y., Y.Y., and N.J.C. co-designed the experiments; H.Y., Y.Y., and D.B. developed analysis methods; H.Y. collected data; H.Y., Y.Y., and D.B., processed and analyzed data; N.J.C. oversaw data analysis, supervised the project, and obtained funding; H.Y., Y.Y., D.B., and N.J.C. interpreted results; H.Y., Y.Y., D.B. and N.J.C wrote the original draft.

## Declare of Competing Interest

The authors declare no competing interest.

## Data and software availability

An archived version of raw and processed datasets and the analysis code will be available through the Johns Hopkins University Data Archive in final submission.

## Material and methods

### Subjects

Adult glass knifefish *Eigenmannia virescens* (length 12 − 15 cm) were purchased from a commercial vendor and housed according to published guidelines [80]. The water temperature and conductivity in fish tank were maintained at 27 ^○^C and 150 – 250 *µ*S/cm. Fish was transferred to the experimental tank at least 12 hours before experiments. Animal care and experimental procedures were approved by the Johns Hopkins Animal Care and Use Committee, and were in accordance with guidelines from the National Research Council and the Society for Neuroscience.

### Experimental apparatus

The experimental apparatus was similar to those in previous work [21, 28, 42]. The setup included a linearly actuated PVC refuge, a 17-gallon rectangular, clear glass tank, a mirror, infrared (IR) lights, and a video camera (Fig. 1A). This system controlled a standardized input trajectory of the refuge, and recorded refuge and fish motion with live video footage.

A 122 cm RGB LED strip (Aclorol, Cincinnati, Ohio, USA) was secured around the tank at about 5 cm below water surface, providing approximately equal amount of illumination within the tank environment. The LED strip was controlled via Arduino Uno (Arduino, Turin, Italy) command running on a computer, providing a maximum illumination of 500 lx. The experimental conditions spanned a wide spectrum on the logarithmic scale. To minimize additional sources of lighting, the experimental space was blocked off with black curtains that spanned from the ceiling down to the floor. The computer monitor was at the minimum brightness level and turned away from the fish tank.

A 30.48 cm × 45.72 cm first surface mirror (FSM first surface mirror LLC, Toledo, OH, USA) was installed at about 45^○^ slanted under the experimental tank, reflecting images of the bottom view of the fish and refuge. Videos were captured by a high-speed camera (pco.1200, PCO AG, Kelheim, Germany) through mirror reflection. Two infrared (IR) lights (CMVision Technologies Inc., Houston, TX, USA) were installed under the experimental tank. These IR lights, invisible to the fish, helped enhance the image quality when recording videos in low light intensities. A 15 cm PVC refuge was attached from the linear actuator to the bottom of the fish tank.

### Stimulus

We designed a pseudo-random periodic sum-of-sines stimulus to control the refuge movements. The input signal was constructed with the following equation, inspired from previous work [41, 42] for system identification of the fish’s frequency-domain tracking responses:

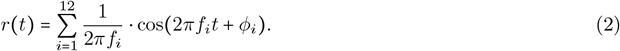

Here, *f*_*i*_ =0.05 × [2, 3, 5, 7, 11, 13, 19, 23, 29, 31, 37, 41] Hz. The phases *ϕ*_*i*_ were randomly chosen to create a unique phase shift for each frequency component. The amplitude of each sinusoidal component was modulated so that the velocity amplitude of each component was kept constant to prevent saturation in tracking high frequencies. Since the sum-of-sines signal contains prime multiples of basis frequency at 0.05 Hz, the period for the stimulus was 20 seconds. With 12 distinct frequency peaks, the time-domain input would be computationally infeasible for the fish to predict [41].

### Experimental Procedure

During experiments, each fish received 5 trials of 9 to 14 illumination stimuli profiles (Table S1). Data was collected with pseudo-randomized trials, where all five fish experienced similar randomization orders, despite slight changes in the numbers of illumination conditions tested between different fish. Before starting each trial, the illumination level was adjusted using an Arduino program. For verification, we put a light meter (LATNEX LM-50KL, Toronto, Ontario, Canada) close to the refuge position near fish tank to measure the illumination level. Each trial lasted for 70 seconds total with a 10-second ramp-up signal at the beginning followed by three 20-second periods of the input signal. Between every two trials, there was a 60-second break with the refuge system paused to prevent fish fatigue.

The linear actuator drove the refuge movement controlled by a custom LabVIEW program [21]. To rule out the effect of fish head direction, we applied the refuge motion trajectory in Equation 2 when the fish head pointed to the right in camera view, and applied the inverted signal of Equation 2 when the fish head pointed to the left. This allows the fish to experience the same tracking task in all trials.

During experiments, we visually inspected real-time tracking plots and identified large events of tracking loss, and then re-ran trials when needed. When it was not possible to keep more than one out of three 20-second repetitions, more follow-up trials were conducted in the original pseudo-randomized order. A video was saved at the end of each trial, capturing the bottom view of refuge and fish at 25 frames per second (FPS). Each video contains 1777 frames (640 pixels × 190 pixels).

Each subject’s data was collected within two consecutive days in up to two experiment sessions, with each illumination level having *at least* five trials (15 repetitions total). Some conditions may have more raw data than others, depending on the difficulty of data collection. The same procedure was standardized and repeated across *N* = 5 subjects.

### Data Preparation

After all data collections, we obtained about 45–52 trial folders for each fish. We then processed videos with custom DeepLabCut [81] machine learning models, MATLAB, and Python OpenCV to obtain 1) the refuge and fish head position, used for fore-aft movement variance analysis and system identification; 2) pixel coordinates along the entire fish body for analysis on fish body bending dynamics. To analyze fish body bending, after obtaining time-series pixel coordinates of the fish head, we shifted each video frame so that the fish head was fixed at a designated “anchor point” at *x* = 220 pixels and *y* = 110 pixels. Another image processing step ensured horizontal alignment of the fish body axis in all frames. This minimized error in tail position analysis due to whole-body rotations. Then, we obtained body-frame position data as shown in Fig. 2A-F. See method details in Supplementary Materials (Fig. S4).

### Data Analysis

#### Managing outliers

The data outliers mainly include: 1) failure of trackers (tracking losses), 2) failure in data collection (i.e. fish left the refuge during experiments), and 3) other trials that fish were in the refuge but did not follow the sum-of-sines trajectory. We first referred to experiment notes and excluded trials in which the fish left the camera view. The time-domain position data (20 seconds each) was plotted in MATLAB for further inspection. Trials with tracking losses were then identified and excluded. We finally identified and excluded other outliers, in which fish were in the refuge but not tracking the sum-of-sines trajectory. For each trial within the same illumination condition, we calculated 1) the sum of squared errors (SSE) for the time-domain position data and 2) the complex multivariate Gaussian distributions of the frequency-domain data, based on Mahalanobis distances. We then established a conservative, balanced metric to exclude outliers, retaining 80% of the data (for more details, see Supplemental Materials: Statistical Methods for Outlier Exclusion).

For body dynamics analysis, we captured pixel coordinates along the entire body up to the tail point. Visual inspection of the original videos and body-frame time-domain data were conducted. This eliminates contaminated trials with poor or hallucinated labels, such as abrupt jumps in body position labels due to DeepLabCut tracking loss. Note that in this analysis, trials that fish did not follow the sum-of-sines trajectory were still included as long as fish whole-body was in camera view.

In some video frames, the tail tip might not captured by the camera. We checked this situation with a simple calculation: if the pixel coordinates of the body exceeded the maximum possible *x* value (*x* = 640), then the trial was excluded. This made sure that all fish body data points were within the camera frame.

#### Fourier Analysis of Fish Tail Movements

When analyzing fish transverse tail movements, we first applied a 2^nd^ order Butterworth low-pass filter with a cutoff frequency of 5 Hz to the time-domain fish body position data sampled at 25 Hz. Then, we extracted the last tail point *b*_11_ in the *y* direction from the centered and aligned fish body position data (see Supplemental Methods: Aligning fish body orientations across video frames). After that, we converted the unit from image pixels to centimeters.

To analyze the frequency components of fish transverse tail movements, we removed the mean from the data to effectively eliminate the DC (0 Hz) component from the signal, as it only represents the constant, non-oscillatory part of the data that is unrelated to the dynamic frequency content of interest. From tail positions, we computed the time-domain fish tail velocities with the forward differences between each two adjacent points, then divided by the sample time 0.04 s. We then calculated the discrete Fourier Transform (DFT) of the time-domain transverse tail positions and velocities using fft function in MATLAB.

To obtain a better qualitative observation of the illuminance-dependence of the above quantities, we applied a modest data smoothing with the MATLAB smooth function, using a window size of three elements. For plotting purposes, we showed the slightly smoothed single-sided DFT amplitude of fish transverse tail positions and velocities, starting from the third data point at 0.10 Hz (Fig. 2G,H, Fig. S2).

#### Frequency Response Function Estimation

Following similar system identification process described in Yang [47], we applied DFT to the time-domain refuge position *r (t*) and fish position *y (t*) across all conditions for *N*= 5 fish, resulting in the refuge position *R [ω]* and fish position *Y [ω]* in the frequency domain.

After that, the frequency response function (FRF) of fish refuge tracking system, *H (jω*), was estimated using empirical transfer function estimate (ETFE) [46]:

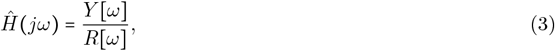

where 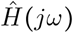 is the estimated fish FRF *H(jω*) using ETFE. Note that this computation is only defined at the 12 frequencies of the sum-of-sine stimulus, since *R [ω]* = 0 at all other frequencies. For the results shown in Fig. 3A-C, we applied the MATLAB smooth function with its default window size of 5 to the FRF gain and phase data across the frequency scale.

#### Frequency-Domain Tracking Error

To examine how the illumination affects refuge tracking performance of fish (Fig. 4I), we computed the frequency domain tracking error *ε(ω*) by the distance between 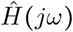 over 12 stimulus frequencies and the perfect tracking point (1, *j*0) on the complex plane, where gain = 1 and phase = 0^○^, i.e.,

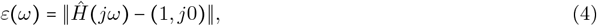

where 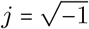.

#### Proposed Computational Model for Sensory Reweighting

We proposed a computational model for sensory reweighting in different illumination conditions based on the assumption that changes in illumination levels would result in reweighting in visual and electrosensory processing.

In our model, we proposed that the “sensory slip” *e (t*) went to two frequency dependent open-loop transfer functions: “*S*_*E*_(*s*)” (open-loop transfer function of electrosense) and “*S*_*V*_ (*s*)” (open-loop transfer function of vision). Two transfer functions were scaled by illumination-dependent weights 1 − *α (λ*) and *α (λ*), respectively, where *λ* is the illumination level in lx. The resulting electrosensory and visual signals were combined and processed into central nervous system *C (s*) and swimming dynamics *P (s*). Thus, the open-loop transfer function was modeled as (1 − *α) P (s) C (s) S*_*E*_ (*s*) + *α P(s) C(s) S*_*V*_(*s*).

Since the FRFs of the closed-loop system were estimated already from experimental data, to open the feedback loop and obtain open-loop FRFs, we followed methods as described in [82]. Using algorithms described in Supplemental Materials, we were able to find illumination-invariant *G*_*V*_ (*s*), *G*_*E*_ (*s*), and a set of *α* values in all illumination scales that minimize the fitting error to open-loop FRFs *G s* computed from experimental data across all illumination conditions.

### Statistical Analysis

All the statistical analysis was performed using custom codes written in R version: 4.3.0, R Core Team, and MATLAB R2024a, MathWorks. The statistical tests performed are: sign test, Mann-Kendall test, Augmented Dickey-Fuller test. The significance level was set to 0.05 for all tests. The experimental and simulation data are provided as either mean plus or minus the standard deviation (*µ*± s.d). or mean plus or minus the standard error of the mean (*µ* ± s.e.m).

## Supplemental Materials

### Supplemental Figures

**Fig. S1.**
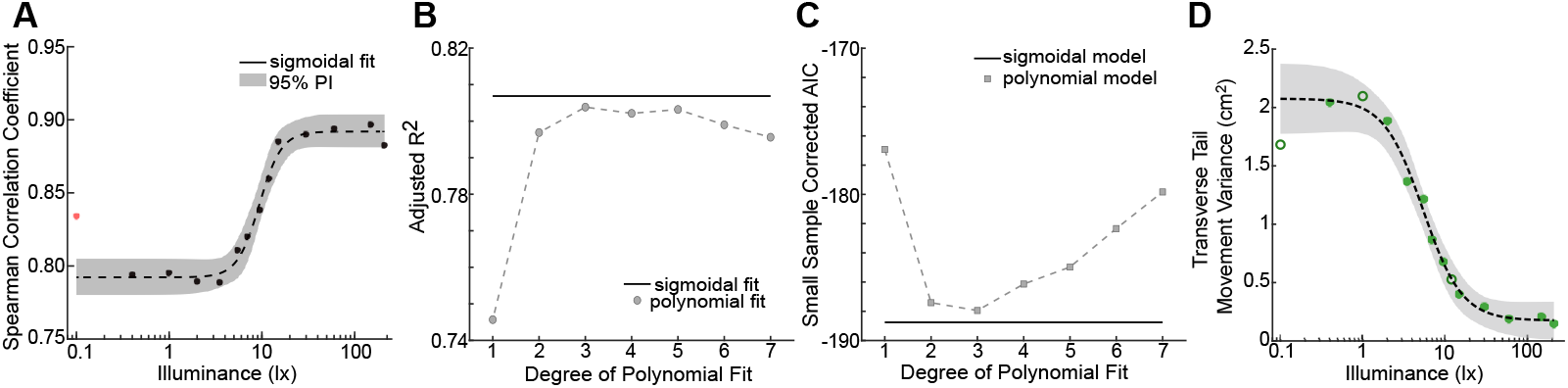
Illumination-mediated switching transition in fish head and tail movements. **(A)** Spearman rank correlation coefficient between the refuge trace and the average fish head movement traces from the same fish from Fig. 1 (fish 1) for different illuminations. High correlation coefficients (> 0.78) observed across all tested illumination levels indicate that the fish consistently followed the refuge. Furthermore, the atypical “S” shape conforms to the switching behavior. The dashed black line and the gray-shaded region denote the sigmoidal fit to the combined data and 95% prediction interval, respectively. The sigmoidal fit to the data was obtained by excluding 0.1 lx data point (red marker) as an outlier. See Table S2 for fitting details. **(B**,**C)** Adjusted *R*^2^ and small sample corrected AIC score substantiate that the sigmoidal model (solid line) is a better fit for the data compared to the polynomial model (markers with dashed line). **(D)** Mean profile for transverse tail movement variance across different illumination for the same fish from Fig. 2I (fish 1). The open circles represent the data corresponding to the neglected outliers from Fig. 2I. The black dashed line and the gray-shaded region denote the sigmoidal fit to the combined data and 95% prediction interval, respectively.

**Fig. S2.**
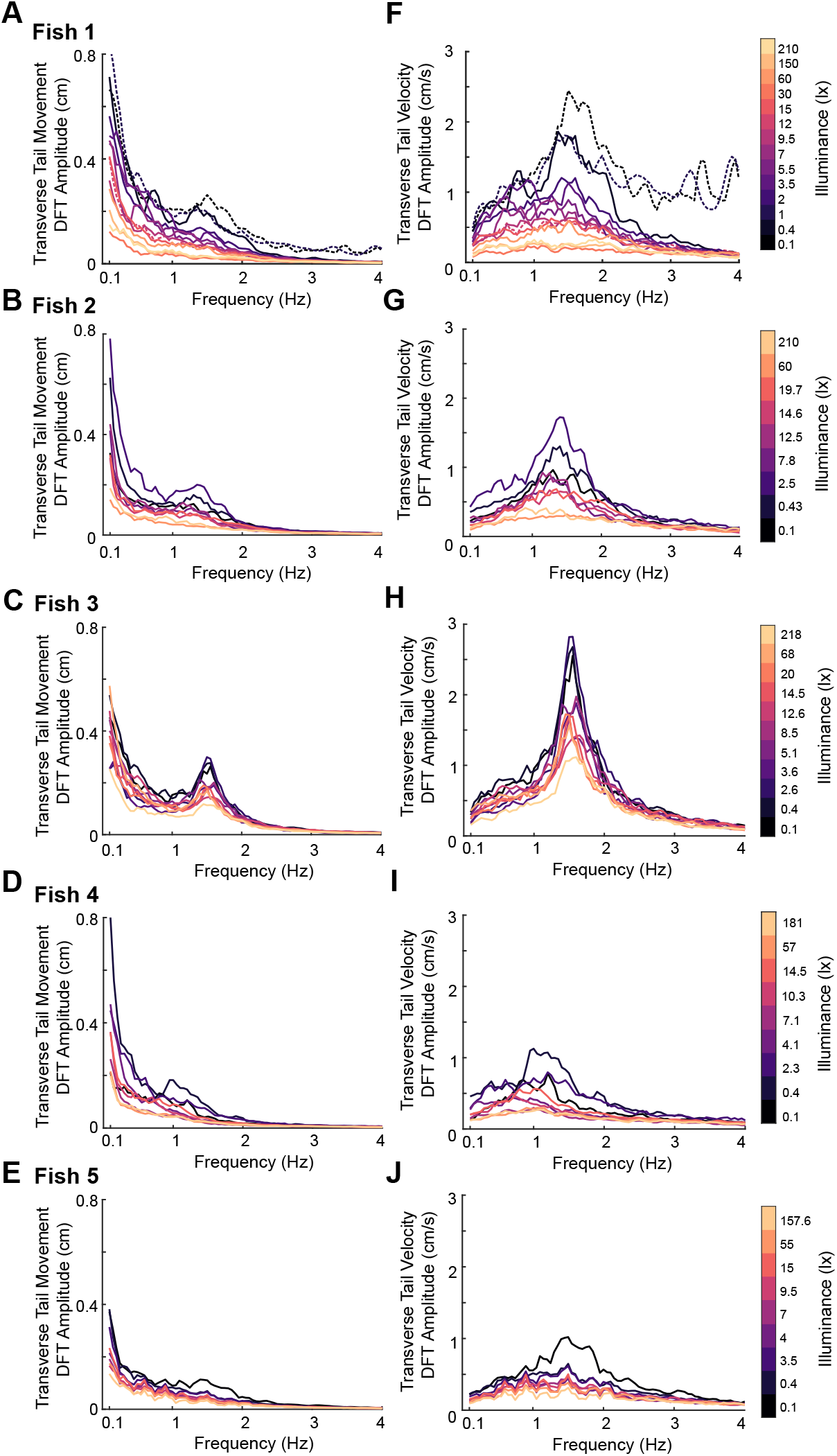
Illumination dependence in transverse tail movement. Average discrete Fourier Transform (DFT) amplitude of transverse tail movement (A–E) and the corresponding velocity (F–J) across trials of each fish (*N=* 5), under different illumination conditions (denoted by different colors). The dashed lines in (A) and (F) represent the data corresponding to the neglected outliers from Fig. 2G,H.

**Fig. S3.**
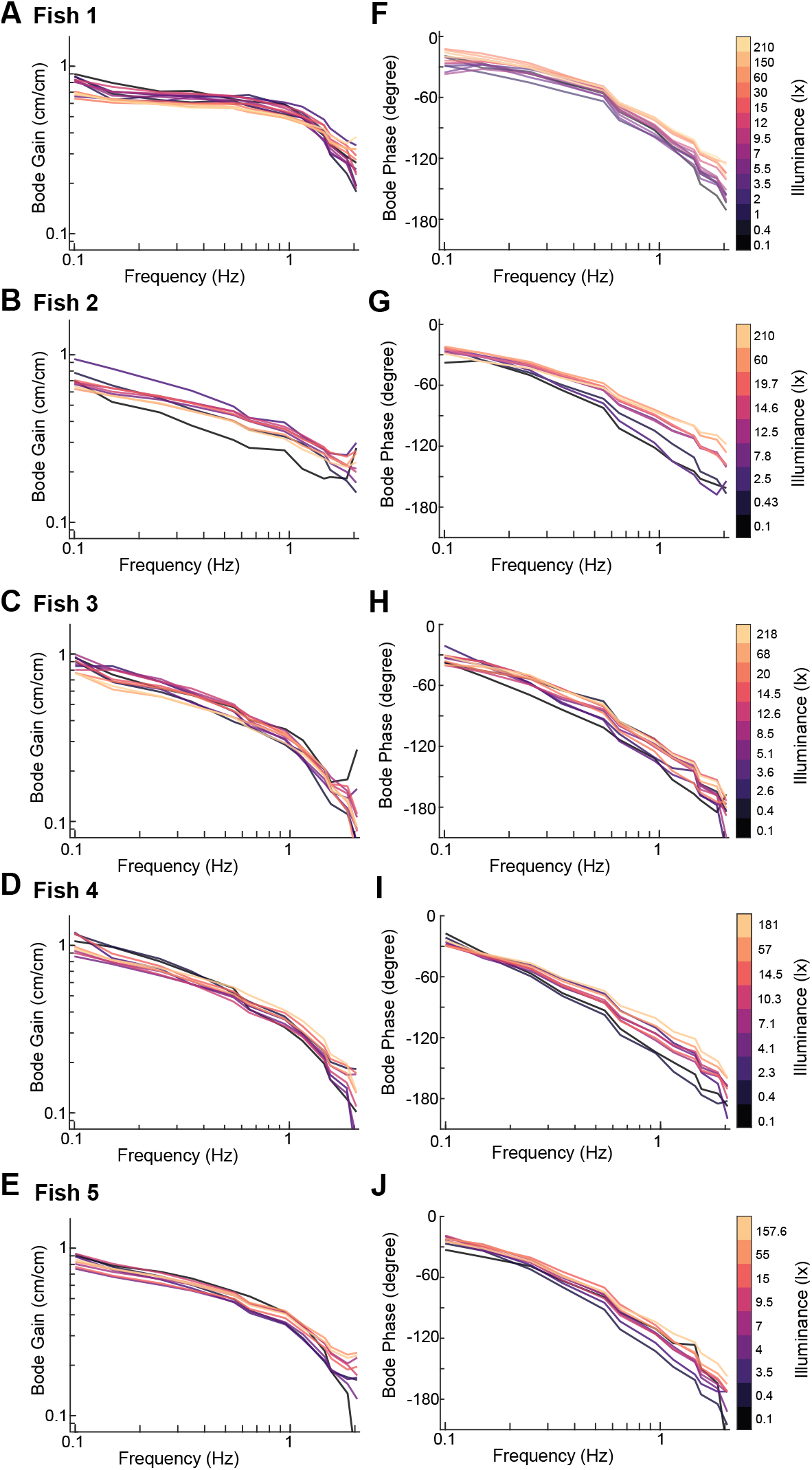
Closed-loop frequency response shows illumination dependent transitions. Bode gain (A–E) and phase (F–J) plots for all individuals (*N* = 5). Different colors represent different illumination levels. The results show that for four out of five fish (except for fish 2), Bode gains tightly distribute over the refuge frequency range (0.1 – 2.05 Hz), whereas for all five fish, Bode phase values are initially tightly grouped but become more dispersed at higher frequencies (> 0.5 Hz).

### Supplemental Tables

**Supplemental Table 1:** The measured illumination levels (lx) of each fish subject. There are slight fluctuations across subjects, due to slight changes in water level of the experimental tank which caused the LED lights to reflect differently.

|  | Fish 1 | Fish 2 | Fish 3 | Fish 4 | Fish 5 |
| --- | --- | --- | --- | --- | --- |
| Condition 1 | 0.1 | 0.1 | 0.1 | 0.1 | 0.1 |
| Condition 2 | 0.4 | 0.4 | 0.4 | 0.4 | 0.4 |
| Condition 3 | 1.0 | 2.5 | 2.6 | 2.3 | 3.5 |
| Condition 4 | 2.0 | 7.8 | 3.6 | 4.1 | 4 |
| Condition 5 | 3.5 | 12.5 | 5.1 | 7.1 | 7 |
| Condition 6 | 5.5 | 14.6 | 8.5 | 10.3 | 9.5 |
| Condition 7 | 7 | 19.7 | 12.6 | 14.5 | 15 |
| Condition 8 | 9.5 | 60 | 14.5 | 57 | 55 |
| Condition 9 | 12 | 210 | 20 | 181 | 157.6 |
| Condition 10 | 15 | — | 68 | — | — |
| Condition 11 | 30 | — | 218 | — | — |
| Condition 12 | 60 | — | — | — | — |
| Condition 13 | 150 | — | — | — | — |
| Condition 14 | 210 | — | — | — | — |

**Supplemental Table 2:** Fitting details for the sigmoidal function, *f* (*x*) = *a*/(1+*e*^−*b*(*x*−*c*)^) +*d* to the data. Data was slightly smoothed by MATLAB smooth function using a moving average filter method with window size of three elements. The fitting parameters, locations of change points in terms of illumination levels (Change Pts, lx), adjusted R^2^ (Adj R^2^), and small-sample corrected Akaike Information Criterion (AICc) are listed below. ΔAdj R^2^ =Adj R^2^ (sigmoidal) − Adj R^2^(best polynomial fit) whereas, ΔAICc =AICc(best polynomial fit) − AICc(sigmoidal). *n* denotes the order of the best fitted polynomial model for that criterion.

| | $a$ | $b$ | $c$ | $d$ | Change Pts (lx) | Adj $R^2$ | AICc | $\Delta\text{Adj } R^2 (n)$ | $\Delta\text{AICc } (n)$ |
| --- | --- | --- | --- | --- | --- | --- | --- | --- | --- |
| Fig. 1I | 2.729 | -2.079 | -0.063 | 0.083 | [0.66, 3.40] | 0.99 | -68.95 | -0.001(4) | 1.08(4) |
| Fig. 1J | 1.691 | -2.015 | 0.192 | 0.093 | [0.95, 5.32] | 0.81 | -140.91 | 0.003(3) | 0.81(3) |
| Fig. 2I | 1.9 | -4.21 | 0.732 | 0.175 | [3.46, 9.46] | 0.99 | -46.10 | $-1.27 \times 10^{-5}(4)$ | 5.62(4) |
| Fig. 2J | 0.606 | -3.183 | 0.845 | 0.453 | [3.46, 12.81] | 0.69 | -196.5 | -0.02(7) | -0.24(3) |
| Fig. 3C | 33.87 | 2.289 | 0.968 | -166.9 | [3.56, 18.17] | 0.86 | 159.38 | 0.003(5) | 2.09(3) |
| Fig. 4F | 0.252 | 2.064 | 0.558 | 0.351 | [1.77, 9.76] | 0.83 | -340.38 | -0.004(6) | 0.64(3) |
| Fig. S1A | 0.100 | 8.374 | 0.973 | 0.792 | [7.09, 12.51] | 0.98 | -122.98 | 0.003(7) | 18.48(3) |

## Supplemental Methods

### Using computational tools to obtain tracked data points from fish video footage

For tracking the fish head positions in time series, we prepared a training dataset of 200 video frames, collected from all subjects under all illumination levels. In each frame, the fish’s head point, *h*_0_, roughly located between the eyes, and one point on the refuge, *s*_0_, were labeled by hand with the DeepLabCut graphic user interface [81] (Fig. S4A).

**Fig. S4.**
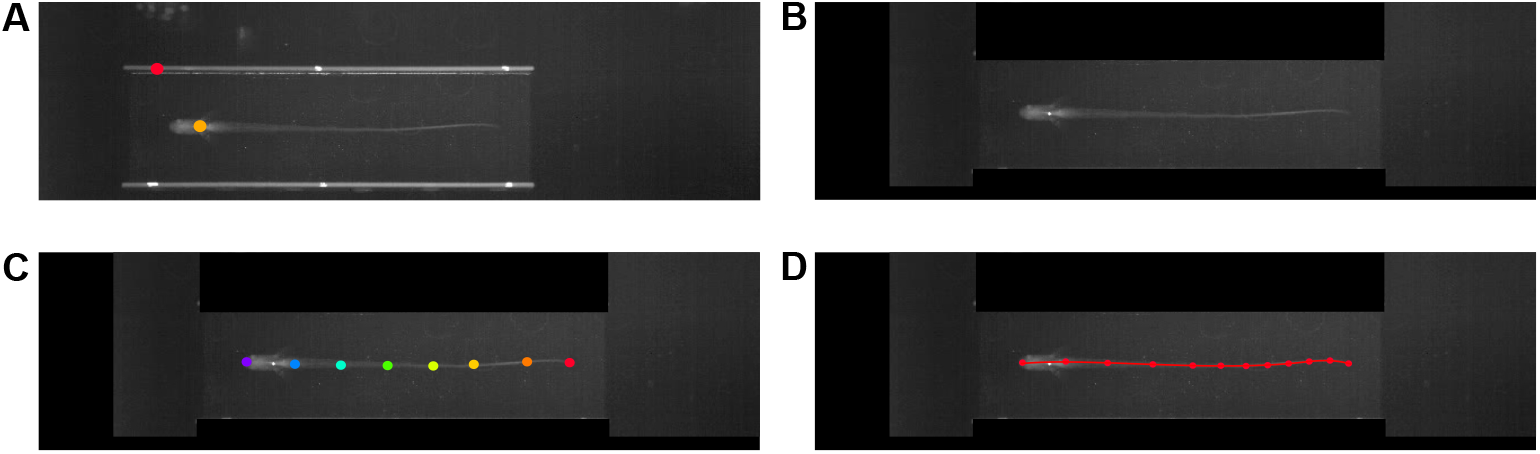
Illustrations of the procedure of tracking fish position and fish body motion. **(A)** The raw video with two points labeled manually: the fish head point and one point on the refuge. **(B)** The pre-processed video with the fish orienting to the left, head pinned at *x* = 220 pixel and *y* = 110 pixel, with the bright rims of the refuge masked out with pixel brightness = 0. **(C)** The DeepLabCut tracked output, with 8 hand-labeled points along the fish body. **(D)** The re-interpolated video output with 12 points, including a red line that connects all the points for visual inspection purposes.

Then, we trained a ResNet-50 machine learning model and analyzed videos to obtain the fish head and refuge positions. The *x* and *y* positions were saved as .csv files in pixel units for each 70-second trial.

To accurately track the entire fish body, we developed an image pre-processing procedure. After obtaining the head position data, we loaded the original videos into a MATLAB script, and then shifted the frame image to pin the fish head point *h*_0_ to the same coordinate: *x* =220 pixels and *y* =110 pixels. To follow the MATLAB image coordinates convention, we standardize the fish orientation to left-pointing. This ensures that the tail always points to the right, where *x* coordinates along the body strictly increase in MATLAB (Fig. S4B).

To minimize chances of image mislabeling in the DeepLabCut model, we masked black rectangles to cover the bright refuge boundaries in each frame, since the fish body would not appear in these regions (Fig. S4B).

From the pre-processed videos, another 200 frames were selected among all trials to train one network with data from all 5 fish. For each frame, we manually picked 8 points that were roughly equally spaced along the fish body. Then, we trained another ResNet-50 model in DeepLabCut. This new model labeled all videos, labeling 8 points along the fish body for each video frame, each at likelihood above 99% (Fig. S4C).

As the manual plotting and fish body-length variability introduced errors in the point spacing, the fish’s body-length curve was re-interpolated again with 12 points. We first bisect the curve, then equally spaced 5 points at the first half and 7 points on the second half. This arrangement helps better capture tail curvature. A Python OpenCV script read the original 8 data points, interpolated a 4^th^ order polynomial to those data points, and segmented the curve into the 12 points mentioned above (Fig. S4D). This ensured that each data point labeled the exact same location on the fish body in every frame; across different fish, this point also labeled the same position with respect to the whole-body length.

### Aligning fish body orientations across video frames

After obtaining the fish head coordinates from the head tip *b*_0_ to the tail end *b*_11_, we aligned the fish’s general body directions across frames. As the fish head positions can be considered rigid (not curving along the body), we fitted a line with the first 3 points from fish head, *b*_0_, *b*_1_, and *b*_2_ and calculated its angle with the horizontal line, *θ*. Then, we rotated all points, *b*_0_ through *b*_11_, around the the head point *b*_1_. This operation ensured consistencies in the body-frame movement analysis, such as the tail-beat motion comparisons shown in Fig. 2A,C,E.

### Statistical Methods for Outlier Exclusion

To exclude outliers within each experimental condition, we implemented a statistical heuristic on both the time-domain head movement data and the frequency-domain tracking responses. The following rules provided a fair metric for data clean-up:

First, if among all trials in a given condition (around 15 trials), a trial’s sum of squared errors (SSE) had a absolute-valued *z*-score greater than 2.2 (98.6% percentile), then it was considered to be an outlier.

Alternatively, we also detected outliers in the frequency domain. With the discrete Fourier transform, each trial would have 12 complex data points for the 12 sum-of-sines frequency components. For each frequency component, we fitted a Gaussian distribution of all trials with the Mahalanobis distance (Fig. S5A). With the 12 *z*-scores per trial, if 1) any one frequency component had *z* > 2.5 (99.4% percentile) or 2) if there were at least three frequency components with *z >*1.5 (93.4% percentile), then the trial was marked as an outlier. Fig. S5B illustrates the outliers identified with the above method, which are categorically different from the rest of the trials. Our data clean-up method ensures that enough data (76% − 100%) remains after excluding outliers, and that the statistical comparisons are fair, since they only compare among trials of the same condition and the same fish. This pipeline preserved accurate and representative data for further processing.

**Fig. S5.**
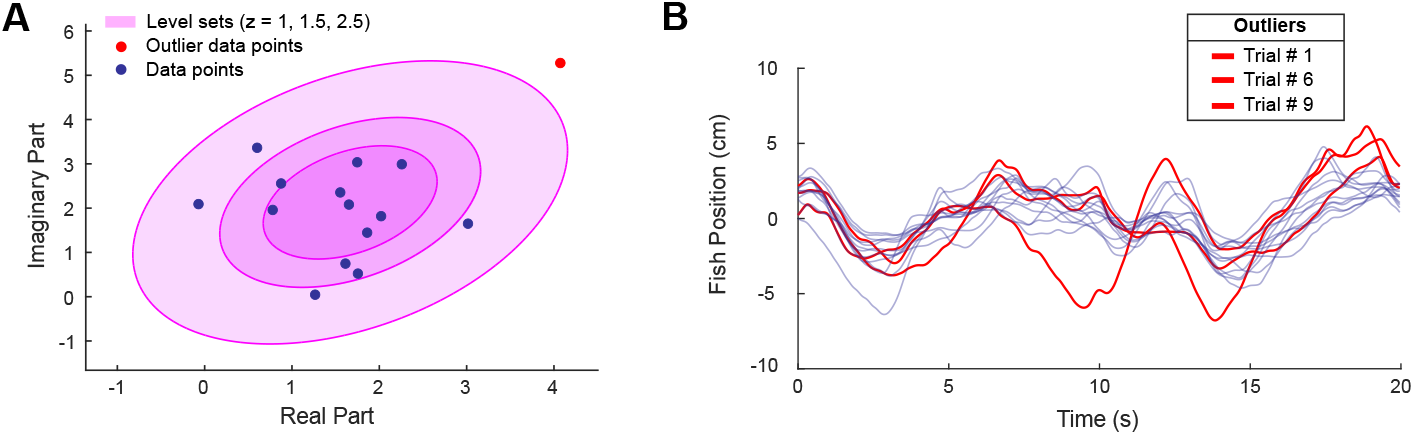
Illustrations of the data clean-up process. **(A)** Within the 18 trials of the same fish and under the same illumination, we plotted the Mahalanobis distances of the tracking responses at a certain frequency peak. Twelve such plots were produced due to 12 frequencies in the sum-of-sines stimulus. Then, we examined the 12 *z*-scores from each trial to identify outliers following the outlier exclusion criteria in Supplemental Methods: Statistical Methods for Outlier Exclusion. **(B)** The time-domain traces of the outliers identified by our metric considering the frequency-domain *z*-scores and time-domain SSE *z*-scores.

### Control-theoretic sensory reweighting model algorithm

We applied an optimization algorithm in MATLAB to obtain illumination-dependent sensory weights in each illumination level, as well as the estimated illumination-invariant non-parametric open-loop transfer functions *G*_*V*_ (*s*) and *G*_*E*_ (*s*) in our proposed sensory reweighting model (see Fig. 4D,G). The estimated non-parametric illumination-invariant open-loop transfer functions, i.e. *G*_*V*_ (*jω*) and *G*_*E*_ (*jω*) as *s* =*jω*, are both 12 complex numbers since there are 12 frequencies in the FRF estimation. We noted them as follows:

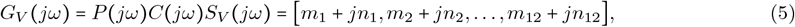

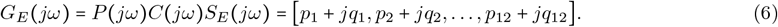

Thus, to determine either *G*_*V*_ (*jω*) or *G*_*E*_ (*jω*) over the frequencies we tested in our experiments, we needed to determine 24 free parameters, i.e., 12 real parameters *m*_*l*_ or *p*_*l*_, and 12 imaginary parameters *jn*_*l*_ or *jq*_*l*_ (*l* 1, 2, …, 12). In our model, we also determined the sensory weights: *α (λ*) for vision (and 1 − *α (λ*) for electrosense) in each of 9 to 14 illumination levels of *λ*. Therefore, depending on the number of illumination levels *λ* we tested on each fish, there were 57 – 62 (24 × 2 +the number of illumination conditions) parameters need to be determined.

Given 500 randomly generated initial conditions for each parameter, we used the fmincon function in MATLAB to find a set of free parameters for each initial condition that minimized a cost function, i.e., the cumulative mean-squared-error (MSE) between model and averaged data at frequencies of interests from all 9 – 14 illumination levels for the specified fish:

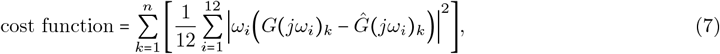

where *k* represents the illumination condition; *n* is the total number of illumination levels (*n* = 9 – 14); *i* represents the frequency of interests (*i* 1, 2, …, 12); terms in [·] is the MSE between averaged data and model in each illumination level; *G (jω*_*i*_) _*k*_ is the estimated open-loop FRF from experimental data at frequency *ω*_*i*_ in the *k*^th^ illumination level 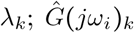 is the fitted open-loop FRF from our sensory reweighting model at frequency *ω*_*i*_ in the *k*^th^ illumination level *λ*_*k*_, and

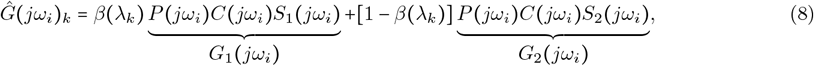

where *β (λ*_*k*_) and 1 − *β(λ*_*k*_), *G*_1_ (*jω*_*i*_) =*P(jω*_*i*_) *C(jω*_*i*_) *S*_1_ (*jω*_*i*_) and *G*_2_ (*jω*_*i*_) =*P (jω*_*i*_*) C (jω*_*i*_) *S*_2_ (*jω*_*i*_) are candidates for sensory weights and open-loop FRF from different sensory pathways, respectively. Due to the fact that fitting result is generally worse at higher frequencies compared with lower frequencies, we multiplied the data and model difference 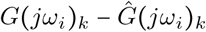 by the frequency *ω*_*i*_ itself, so that the cost function put higher weights on errors at higher frequencies.

Although the form of Equation 8 is similar to that of the sensory reweighting model in Equation 1, we need to further determine whether *α* (*λ*_*k*_) = *β* (*λ*_*k*_) or *α* (*λ*_*k*_) = 1 − *β* (*λ*_*k*_), and which one of *G*_1_ (*jω*) and *G*_2_ (*jω*) is *G*_*E*_(*jω*) and which one is *G*_*V*_ (*jω*) since the optimization algorithm itself treated each sensory pathway equally. only giving us candidates for sensory weights and open-loop FRF from different sensory pathways. Therefore, we used another algorithm to determine each item by checking the trend of *β* (*λ*_*k*_). In the lowest illumination level 0.1 lx, it makes sense that fish is “electrosense dominated”, i.e., they put lower weights on vision since they cannot see very well and put higher weights on electrosense. Thus, *α* at 0.1 lx should be relatively small and 1 − *α* should be relatively large. As illumination increased, the weight *α* in visual pathway should be increasing and *α*(*λ*_1_) < *α*(*λ*_*n*_). Therefore, if *β*(*λ*_*k*_) obtained from the optimization algorithm satisfies *β*(*λ*_1_) < *β*(*λ*_*n*_), then

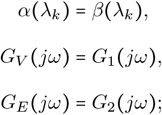

otherwise,

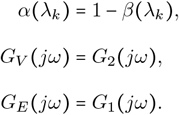

Note that sensory weights for vision *α* and that for electrosense 1 − *α* depend on illumination levels *λ*_*k*_, while visual and electrosensory pathways *G*_*V*_ (*jω*) and *G*_*E*_(*jω*) are illumination-invariant. In the end, among all 500 best fitted parameter sets for each initial condition, we selected the set of parameters that gave the least fitting error as the final parameters of fitting for each fish.

### Comparing the bootstrapped experimental data with the sensory reweighting model

To better quantify the fitting quality, we conducted bootstrapping analysis for each fish (*N* = 5). The procedure is as follows: in each illumination condition, we created a sampled dataset by sampling trials from the actual dataset with replacements. After repeating this step for all illumination conditions, we created a bootstrapped dataset. Following those steps, we generated 100 bootstrapped datasets for each fish (*N* = 5). For each bootstrapped dataset, we applied sensory reweighting control model algorithm to obtain the corresponding *G*_*E*_ (*jω*) and *G*_*V*_ (*jω*) and a set of illumination-dependent visual weight *α*. Thus, for each fish, we finally obtained 100 bootstrapped datasets and their corresponding fitted parameters in each illumination. All code for this bootstrap analysis was run in MATLAB R2024a.

